# Quantitative profiling of pseudouridylation landscape in the human transcriptome

**DOI:** 10.1101/2022.10.25.513650

**Authors:** Meiling Zhang, Zhe Jiang, Yichen Ma, Wenqing Liu, Yuan Zhuang, Bo Lu, Kai Li, Chengqi Yi

## Abstract

Pseudouridine (Ψ) is an abundant post-transcriptional RNA modification in ncRNA and mRNA. However, transcriptome-wide measurement of individual Ψ sites remains unaddressed. Here, we develop “PRAISE”, via selective chemical labeling of Ψ by bisulfite to induce nucleotide deletion signature during reverse transcription, to realize quantitative assessment of the Ψ landscape in the human transcriptome. Unlike traditional RNA/DNA bisulfite treatment, our approach is based on quaternary base mapping and revealed a ~10% median modification level for 2,714 confident Ψ sites in HEK293T cells. By perturbing pseudouridine synthases, we obtained differential mRNA targets of PUS1, PUS7 and TRUB1, with TRUB1 mRNA targets showing the highest modification stoichiometry. In addition, we identified and quantified known and novel Ψ sites in mitochondrial mRNA, catalyzed by a mitochondria-localized isoform of PUS1. Collectively, we provide a sensitive and convenient method to measure transcriptome-wide Ψ; we envision this quantitative approach would facilitate emerging efforts to elucidate the function and mechanism of mRNA pseudouridylation.

## Introduction

More than 170 different post-transcriptional RNA modifications have been identified to date^1^. Pseudouridine (Ψ), known as the “fifth nucleotide” of RNA, is the most abundant modification and occurs in all three domains of life^2^. It was first discovered in 1951 and is widespread in ribosomal RNA (rRNA), small nuclear RNA (snRNA) and transfer RNA (tRNA)^3–5^. In messenger RNA (mRNA) of mammalian cells, it is also abundant and represents the second most abundant internal mRNA as measured by quantitative mass spectrometry^6^. By now, Ψ has been found to impact various biological processes including translation, splicing, RNA stability, protein–RNA interaction and response to cellular environments^7,8^. Recently, its importance is further exemplified by the success of COVID-19 mRNA vaccine, which is modified by Ψ and its methylation derivatives^9,10^.

Ψ is catalyzed by snoRNA-dependent ribonucleoprotein or guide RNA independent, stand-alone pseudouridine synthases (PUS)^4^. There are 13 PUS members that belong to five families (TruA, TruB, TruD, RluA and PUS10) in eukaryotes. Ψ synthases have been found to affect cellular activities as well as to contribute to pathological conditions. Within TruD family, pseudouridine synthase 7 (PUS7) mediated pseudouridylation can affect cap-dependent protein translation and contribute to stem cell dysfunctions by altering the properties of mTOG tRNA derived fragments (tRFs)^11,12^. In addition, PUS7-mediated Ψ modification inhibits codon-specific translation of tRNAs in glioblastoma and catalytic inhibition of PUS7 can reduce tumorigenesis^13^. PUS1, PUS7 and RPUSD4 have been found to modify human pre-mRNA co-transcriptionally and modulate alternative pre-mRNA processing^14^. Some functions of PUS are independent of their activity, for instance, pseudouridine synthase 10 (PUS10) play a role in miRNA biogenesis by promoting pri-miRNA processing, which is independent of its catalytic activity^15^. TRUB1 is also reported to regulate the maturation step of miRNA let-7 in an enzyme activity-independent manner^16^. Aberrant expression or catalytic activity of Ψ synthases have been linked to various human diseases, including mitochondrial myopathies and intellectual disability^7^. Mitochondrial myopathy and sideroblastic anemia (MLASA) have been associated with a missense mutation in pseudouridine synthase 1 (PUS1), an enzyme belonging to the TruA family^17^. A homozygous truncating mutation in PUS3 (also TruA family) contributes to intellectual disability^18^. Inactivating mutations in DKC1 encoding an RNA-dependent TruB member were found to be associated with X-linked dyskeratosis congenita^19^.

Our study and understanding of RNA modifications is greatly facilitated by advances in modification detection approaches, especially high-throughput sequencing methods to profile the transcriptome-wide modification landscape. Because Ψ is a “silent” modification during reverse transcription and by mass spectrometry, a selective chemical labeling strategy utilizing N-cyclohexyl-N’-β-(4-methylmorpholinium) ethylcarbodiimide p-tosylate (CMCT) has been developed to identify Ψ in a primer-extension assay^20^. This chemistry has also been coupled to high-throughput sequencing to enable transcriptome-wide detection of Ψ in yeast and human cells^6,21–23^. Other than CMCT, additional chemistry has been proposed to realize selective reactions to Ψ but not regular U in RNA^24–28^. More recently, nanopore-based direct RNA sequencing has been developed to profile RNA modifications including Ψ^29–31^.

Despite the progresses, measurement of Ψ stoichiometry in the transcriptome remains unaddressed. As a matter of fact, identification of Ψ sites in the transcriptome is still problematic at the moment. While the four NGS-based methods first enabled transcriptome-wide profiling of Ψ, the limited overlap of modification sites between the studies suggest that their accuracy and specificity remain to be improved^32^. More specifically, the CMC-based methods suffer from false positives and false negatives, which originate from RNA structure-induced RT stops and partial CMC labeling. Nanopore sequencing methods were designed to directly detect RNA modifications based on base-calling error, which are bioinformatically demanding and highly sensitive to the abundance of the transcripts as well as modification level. Hence, a quantitative, accurate and sensitive method for global Ψ detection is urgently needed.

In this study, we develop a transcriptome-wide method to quantitatively detect Ψ at single-base resolution. This method relies on bisulfite-induced deletion signature during reverse transcription, thus enabling quantitative pseudouridine assessment via bisulfite/sulfite treatment (named “PRAISE”). PRAISE is based on quaternary reads alignment and thus accurately measures Ψ stoichiometry in spike-in RNA and as well as rRNA. In the transcriptome of HEK293T cells, we were able to identify 2,714 confident Ψ sites, showing a ~10% median modification level. Combining PRAISE with pseudouridine synthases perturbations, we assigned differential Ψ sites to PUS1, PUS7 and TRUB1 enzymes. Moreover, we identified known and novel Ψ sites in mitochondrial mRNA, which are catalyzed by a mitochondria-localized isoform of PUS1. Collectively, our approach reveals the quantitative landscape of Ψ in the human transcriptome and provides a reliable and sensitive method for the elucidation of biogenesis and function of mRNA pseudouridylation.

## RESULTS

### Bisulfite-Ψ adduct induces deletion during reverse transcription

We sought to identify a small molecule that could effectively distinguish Ψ from regular U. Interestingly, literatures from more than half a century ago reported that pseudouridine can be irreversibly modified by bisulfite to yield two ring-opening monoadducts^33,34^ (Fig.1a). A later study found that ring-opened abasic site structures in DNA can block translesional synthesis and lead to deletions during DNA synthesis^35^. We thus hypothesized that Ψ in RNA could be labeled by the bisulfite reaction and subsequently generate nucleotide deletions during reverse transcription.

**Figure 1.**
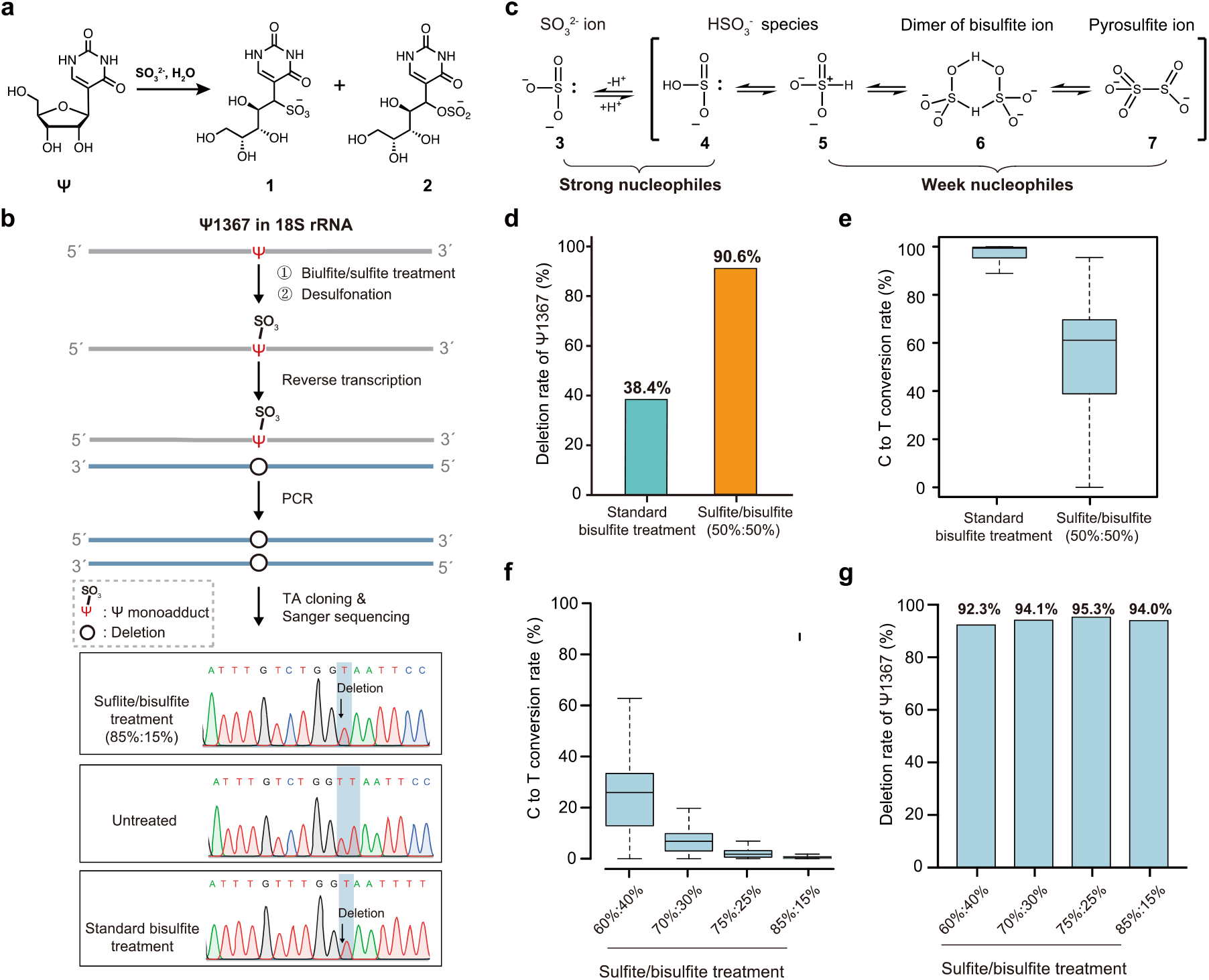
Bisulfite-Ψ adduct induces nucleotide deletion during reverse transcription. (a) The reaction of bisulfite with Ψ to yield two ring-opened, mono-bisulfite adducts **1** (S-adduct) and **2** (O-adduct). (b) Scheme of bisulfite reaction with Ψ. HEK293T total RNA and detection for Ψ1367. The bisulfite reaction contains two sequential steps, which are bisulfite/sulfite treatment and desulfonation. RNA was then reverse transcribed followed by targeted PCR, TA cloning and sanger sequencing of individual clones for Ψ1367 in 18S rRNA. (c) Structure of sulfite and bisulfite ions and transformations among them. In bisulfite solution (pH=5.1), sulfite ion **3** (SO_3_^2−^) is in equilibrium with bisulfite species (HSO3^−^) **4** and **5**, which are two tautomers. At high concentration of bisulfite (>10^−2^ M), these two tautomers interact by hydrogen-bonding and convert to bisulfite ion dimer **6**, which is in turn in equilibrium with the pyrosulfite ion **7** (S_2_O_5_^2−^). (d) The deletion rate of Ψ1367 site in 18s rRNA under standard bisulfite condition and 50%sulfite/50%bisulfite condition. (e) The conversion rate of C-to-T under standard bisulfite condition (cyan) and 50%sulfite/50%bisulfite condition (orange). (f) The conversion rate of C-to-T under different proportion of sulfite and bisulfite. The ratio of the sulfite reagent ranges from 60% to 85%, and the conversion rate significantly decreases with the increase of sulfite ratio. (g) The deletion rate of Ψ1367 site in 18s rRNA under different sulfite/bisulfite conditions. The ratio of sulfite reagent ranges from 60% to 85%, and the deletion rate remained stable with the increase of sulfite ratio.

Since bisulfite treatment has been adopted for m^5^C detection in RNA^36,37^, we applied the literature reaction (3.8 M bisulfite solution, pH=5.1) to total RNA of HEK293T cells and analyzed the signal of Ψ1367, a highly modified Ψ site in 18S rRNA. We were able to find a significant deletion signature for the site, showing a 38% deletion rate (Fig.1b,d). The result is consistent with recent papers reporting optimized reaction conditions for transcriptome-wide m^5^C detection, in which Ψ is simultaneously detected as a side product^28,36^. While the data clearly shows the potential of bisulfite in Ψ detection, the chemistry suffers from two fundamental limitations. First, Ψ stoichiometry is severely underestimated: we only observed a 38% deletion signal for a site that is modified to greater than 95% level. Second, the current bisulfite condition converted all cytosines to uridines. Such reduced sequence complexity is expected to lead to low mapping rates and inaccurate Ψ identification particularly within CU-enriched regions.

To tackle the first challenge and improve the reaction efficiency, we looked into the chemical mechanism of the reaction. While bisulfite (HSO_3_^−^) is believed to act as the effective nucleophile, literature in the 1970s has clearly documented that within the high concentration of sodium bisulfite solution, the predominant component is bisulfite ion dimer and pyrosulfite ion (S_2_O_5_^2−^)^38^, which is inactive towards Ψ (Fig.1c). Hence, we speculated that by increasing the proportion of sulfite and decreasing that of bisulfite, the effective nucleophile concentration can be improved while the total level of HSO_3_^−^ and SO_3_^2−^ ions remains unchanged. The traditional bisulfite reagent (pH=5.1) is equivalent to a molar mixture of 95% bisulfite and 5% sulfite (bisulfite:sulfite=19:1). We found that when the molar proportion of sulfite was increased to 50%, the deletion rate of Ψ1367 was increased to 90% (Fig.1d, Supplementary Fig.1a). Among the deletion signals, 1-bp deletion accounts for 90%, and 2-bp deletion accounts for 10%. Hence, the efficiency of bisulfite-Ψ addition reaction can be greatly improved by increasing the concentration of the effective nucleophile.

We then set out to solve the C-to-U conversion problem. Under the improved condition for deletion detection (bisulfite:sulfite=1:1), we still observed a high C-to-U conversion rate (~60%) (Fig.1e). Based on the mechanism of bisulfite reaction, the deamination of cytosine prefers weakly acidic conditions while the addition of bisulfite to Ψ could occur under relatively basic conditions (Supplementary Fig.1b,c). We thus reasoned that we should be able to minimize C-to-U conversion by further increasing the portion of sulfite ion, which increases not only the pH value of the reaction condition but also the effective nucleophile concentration. We tested a range of sulfite proportion (from 60% to 85%, corresponding to a pH value from 7.0 to 7.8), and found that the conversion rate significantly decreased with higher proportion of sulfite (Fig.1f). Notably, under the 85% sulfite condition, an extremely low C-to-U conversion rate (< 0.5%) was observed, without compromising the deletion signal of Ψ (Fig.1g). We did not try higher proportion of sulfite because a strong alkaline environment may lead to severe RNA degradation and hence affect library construction efficiency.

Lastly, we optimized some of the key steps of the bisulfite reaction, including the incubation temperature, incubation time and desulfonation time, as well as the choice of RTases (Supplementary Fig.1d,e). We confirmed that the most optimal condition was saturated solution containing 85% sulfite and 15% bisulfite for 5 h at 70°C, and that desulfonation condition was 1 M Tris pH 9.0 for 0.5 h at 75°C (see methods). Fragment analysis suggested that the bisulfite reaction caused visible but acceptable degradation of RNA molecules (Supplementary Fig.1f). We further systematically compared the performance of several commercially available RTases (including SuperScript II, SuperScript III, SuperScript IV, Maxima H minus, Revert Aid, M-MLV, AMV, ProtoScript II, Recombinant HIV and HiScript III) under their recommended RT conditions. We found that Maxima H minus, which is used in scRNA-seq (by Smart-seq3 and et al.)^39,40^, demonstrated excellent RT efficiency and high deletion frequency at the sites of Ψ-bisulfite adduct (Supplementary Fig. 1 g). Collectively, we identified an optimized bisulfite condition that minimizes C-to-U conversion while maximizes Ψ detection based on chemical-induced deletion signature during RT.

### A customized bioinformatics pipeline of PRAISE

We then coupled the labeling reaction with next-generation sequencing (NGS) to develop the “PRAISE” technology for global identification and quantification of individual Ψ sites (Fig.2a). While a deletion signal is relatively straightforward to analyze in Sanger sequencing, there exists several challenges during NGS data analysis. First, a deletion signal induced by the Ψ-bisulfite adduct is equivalent to a gap located internally in the sequencing reads. The default setting of existing mapping software is to remove reads with gaps, especially those with multiple gaps. Thus, current mapping strategies are anticipated to cause underestimation of Ψ stoichiometry or even false negatives in Ψ identification, particularly for RNA sequences with dense Ψ sites. Another problem is that when a gap is present within the seed region during read mapping, the gap will be soft-clipped, eventually leading to loss of genuine modification sites. In order to deal with these challenges, we proposed a bioinformatics pipeline composed of two key procedures. In the first step, we chose hisat2^41^ and used the “very sensitive” parameter to prevent over-filtering of sequencing reads. In the second step, we carried out local alignment using a tailored scoring matrix and filter to obtain accurate mapping results (see methods). In case there is a gap in the seed region, we retain the gap and extend the reference sequence to fit results with deletion signals. After incorporating these changes, we were able to achieve more accurate deletion signals and hence Ψ profiling. For instance, there are four highly modified Ψ sites in the region 3,665-3,720 of 28S rRNA^42^. Using the default alignment parameters, the modification rate was approximately ~30%, representing significant underestimation of modification level. After adopting our in-house made alignment method, we observed deletion rates of about 90% for the four Ψ sites, while the flanking bases exhibited very low deletion rate (< 1%) (Fig.2b). Thus, we believe that the customized pipeline is suitable for PRAISE-enabled Ψ identification.

**Figure 2.**
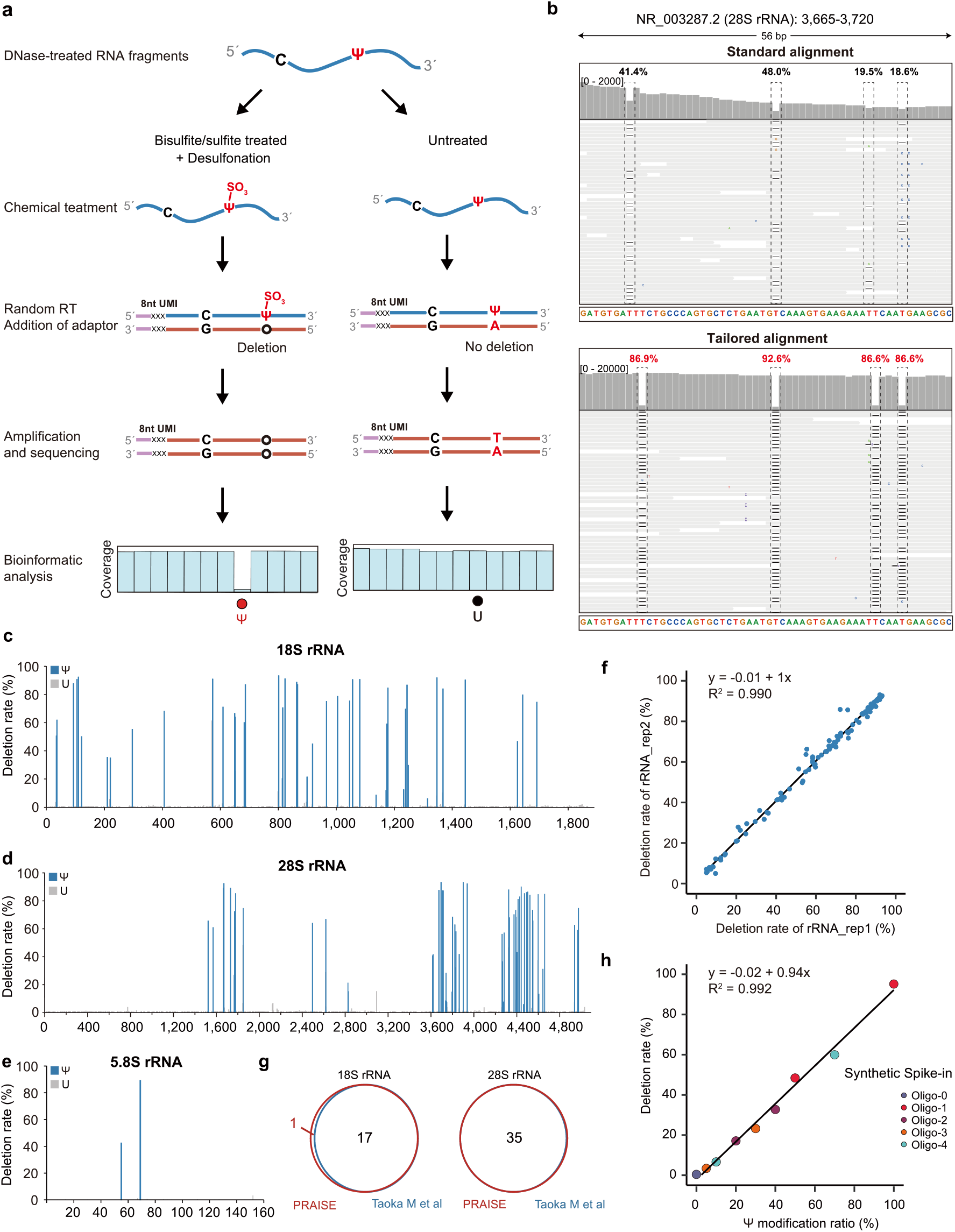
PRAISE accurately measures Ψ stoichiometry in rRNA and synthetic spike-in oligos. (a) Schematic illustration of PRAISE. DNase I treated RNA is first fragmentated to ~150 nt, and treated with the optimized bisulfite/sulfite conditions. “Untreated” sample is set as a negative control. After library construction, Ψ sites are identified as deletion signals during sequencing. An 8 nt unique molecular identifier (UMI) is added through the reverse-transcription step to remove potential PCR duplications, further improving the confidence of quantitative Ψ identification. (b) Representative IGV results for dense Ψ sites within region 3,665-3,720 of 28S rRNA. The NGS data after bisulfite/sulfite treatment is mapped by default (upper panel) or customized strategies (lower panel), respectively. Four highly modified Ψ sites, including Ψ3672-3674, Ψ3694, Ψ3708-3709, and Ψ3713 are shown by the dotted lines. (c) PRAISE reliably detects Ψs in 18S rRNA. The deletion rates in one replicate are shown. (d) PRAISE reliably detects Ψs in 28S rRNA. Many Ψ sites tend to densely locate in a short region, for instance two regions near the 3’ end of 28S rRNA. (e) PRAISE reliably detects the 2 Ψs in 5.8S rRNA. (f) Deletion rates between the two biological replicates, showing a high quantitative reproducibility of PRAISE. (g) PRAISE quantifies Ψ stoichiometry of 5 synthetic spike-in RNA. Scatterplot comparing deletion rates measured by PRAISE (y axis) with Ψ stoichiometries as expected (x axis). The Pearson correlation coefficient (R^2^) and the trend line equation are denoted.

### PRAISE sensitively measures Ψ stoichiometry in rRNA

We then applied the bioinformatics approach to detect Ψ modification in rRNA. In total, PRAISE identified 45, 62 and 2 Ψ sites in 18S, 28S and 5.8S rRNA, respectively (Fig 2c,d,e, Supplementary table 1). PRAISE also demonstrates excellent repeatability for Ψ ratio measurement between two technical replicates (Pearson correlation: R^2^= 0.990) (Figure 2f). Since rRNA has been extensively characterized for various post-transcriptional modifications, we also compared the sequencing results by PRAISE with orthogonal mass spectrometry data^42^. We classified all Ψ sites into two scenarios: Ψ within single U context or within consecutive Us. For both situations, we observed a high consistency between the two methods: for the 17, 35 and 2 Ψ sites residing in single U context in 18S, 28S and 5.8S rRNA, they were all detected unambiguously (Figure 2g); for the 26 and 26 known Ψ sites residing in consecutive U context in 18S and 28S rRNA, PRAISE can’t tell the exact modification position but is capable to report the modification within the consecutive U sequence. For instance, region 1847-1850 of 28S rRNA has four consecutive Us; while PRAISE can’t report the exact site of modification, a closer examination of the IGV view clearly suggests the presence of two discontinuous Ψ sites within the region (Supplementary Fig.2a). Besides reliably detecting the known Ψ sites (with reference to the MS and database results^43^), PRAISE also identified three additional Ψ sites in rRNA, which are Ψ1172 in 18S rRNA, and one Ψ site within U1314-1315 region of 18S rRNA and the U4094-4095 region of 28S rRNA respectively. Notably, our previous clickable CMC-based CeU-seq technology also captured the latter two modification sites, providing orthogonal evidence to the presence of Ψ1315 in 18S rRNA and Ψ4095 in 28S rRNA^6^. To further validate the novel Ψ1172 site in 18S rRNA, we knocked down the DKC1 enzyme, which is responsible for rRNA pseudouridylation (Supplementary Fig.2b,c,d). Indeed, Ψ1172 showed a significantly decreased deletion rate in the DKC1 KD samples, as well as the other two Ψ sites (Figure 2e). In addition, derivatives of Ψ modification, including hypermodified m^1^acp^3^Ψ1248 in 18S rRNA and Ψm3797 in 28S rRNA, could also be identified at high confidence (Supplementary Fig.2f). Moreover, we investigated the performance of PRAISE towards other rRNA modifications, such as 2’-O-methylation (Um, Cm, Gm, Am), m^6^A, m^6^_2_A, m^5^C or ac^4^C. We did not find any of these sites showing a deletion rate greater than 2%, demonstrating the high specificity of PRAISE (Supplementary Fig.2g,h).

To further assess the quantitative capability of PRAISE, we chemically synthesized 5 spike-in RNA oligos containing a single Ψ site respectively, mixed the fully modified RNA with the matched, unmodified RNA to ratios ranging from 0% to 100% (0%, 5%, 10%, 20%, 30%, 40%, 50%, 70%, 100%) and measured the deletion signal in PRAISE. We achieved excellent quantitative agreement (Pearson correlation: R^2^= 0.992) between the expected Ψ values and the experimentally determined Ψ levels, even at the 5% modification level (Fig.2h). Taken together, sequencing results from both rRNA and spike-in oligos showed that PRAISE is capable of quantitatively detecting Ψ sites.

### A quantitative Ψ landscape in the human transcriptome

Having validated the unbiased and quantitative ability of PRAISE, we next characterized the transcriptome-wide Ψ modification sites and stoichiometry. We were able to identify 2,960 and 2,971 Ψ sites in the two biological replicates respectively, with 2,174 shared Ψ sites (~73%), thus demonstrating high reproducibility of PRAISE (Fig.3a, Supplementary table 2). To evaluate the reliability of our method, we employed a recently established in vitro transcribed RNA (IVT RNA) library from HEK293T transcriptome as a negative control^44^. We first confirmed that ~80% genes were recovered in the IVT library, and the read coverages across transcripts were similar in IVT and PRAISE libraries (Supplementary Fig.3a,b). We could only identify 22 and 14 Ψ sites in two technical replicates of IVT RNA respectively, with 5 shared Ψ sites (Supplementary Fig.3c). This observation also suggested the high confidence of PRAISE. Moreover, we compared the 5 false-positive sites from IVT RNA with the 2,174 Ψ sites identified from cellular RNA, and found that only 3 Ψ sites overlapped (Fig.3b). Therefore, the Ψ landscape obtained via PRAISE is highly reliable. We also showed the sequencing data for one representative example, showing ~90% deletion rate in the replicates of celluar RNA while <1% deletion rate in the IVT replicates (Fig.3c).

**Figure 3.**
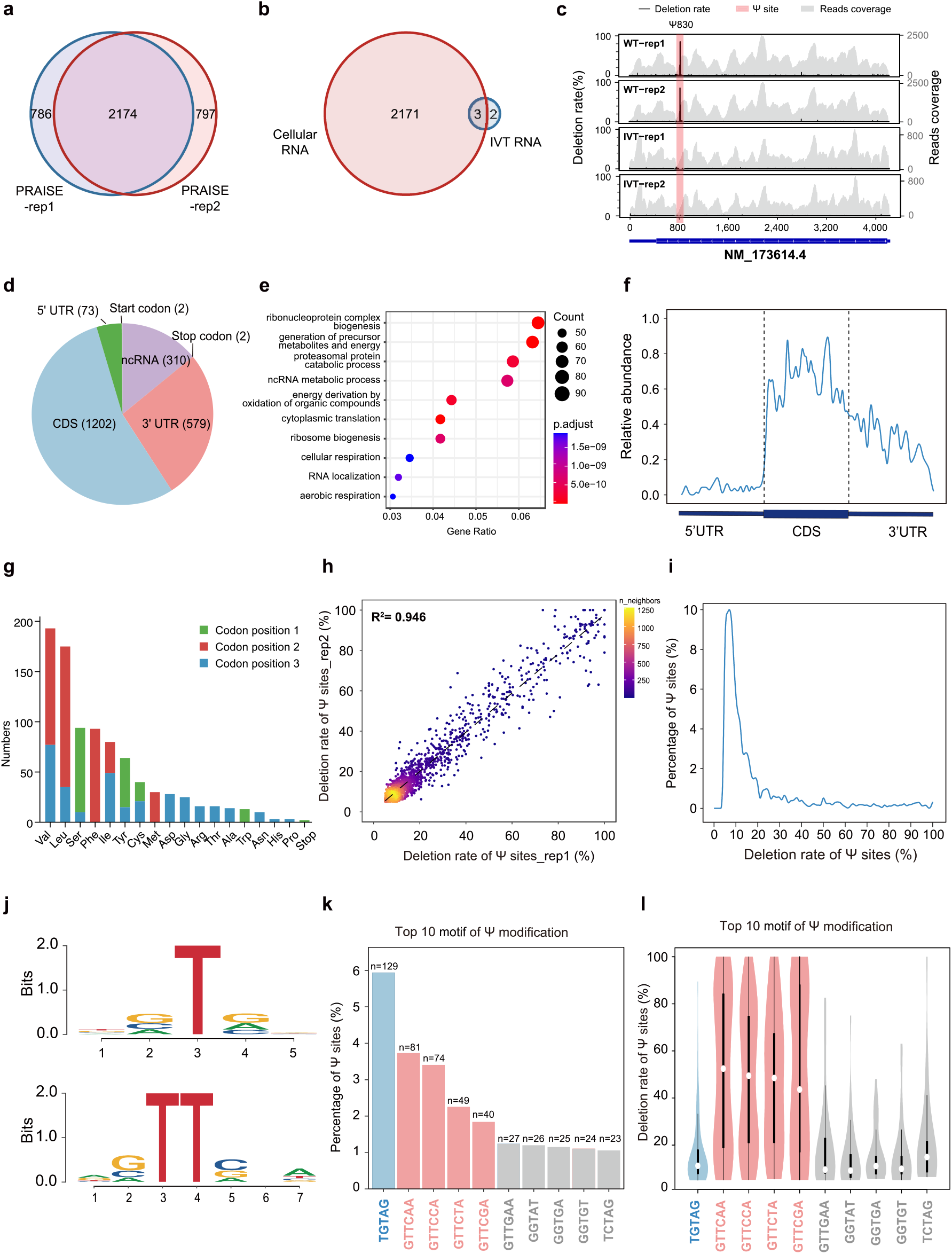
The quantitative landscape of Ψ in the human transcriptome. (a) Venn diagram shows the overlap of Ψ sites in the two biological replicates. (b) Venn diagram shows the overlap of Ψ sites in the cellular RNA library (2,174 Ψ sites) and the modification-free, IVT RNA library (5 Ψ sites). (c) Representative views of one Ψ site on the transcript of NM_173614.4 (NOMO2). Deletion rate (left y-axis) and read counts (right y-axis) are both shown in PRAISE-rep1, PRAISE-rep2, IVT-rep1 and IVT-rep2 samples. Gray background color denotes reads coverage. Ψ site is indicated with a red background. (d) GO enrichment analysis (BP) for the 2,174 transcriptome-wide Ψ sites in a. (e) Metagene profiles showing the distribution of Ψ sites in human mRNA. (f) Pie chart showing the proportion of Ψ sites in mRNA and ncRNA. (g) The counts of Ψ sites in different codons as well as positions are shown. (h) Deletion rates of all mRNA Ψ sites between the two biological replicates, showing a high correlation. The color gradient denotes the density of overlapping modification sites. (i) Curve graph showing the proportion of Ψ sites with different modification levels. (j) Motif analysis of identified Ψ sites within “U” (upper panel) and “UU” contexts (lower panel). (k) The proportion of top 10 sequence contexts containing Ψ sites. The blue and red color denote the TGTAG and GTTCNA motif, respectively. Grey color denote the remaining motifs. (l) The deletion rate of Ψ sites within the top 10 motifs.

Among the 2,174 shared modification sites, 1,864 and 310 sites are located in mRNA and ncRNA (excluding rRNA) of HEK293T cells respectively (Fig.3d). To determine if Ψ modification is linked to specific cellular processes, we performed Gene Ontology (GO) term enrichment analyses of Ψ-containing transcripts. The GO analysis of BP (Biological Process) found highly significant (*p* < 2.5e-07) enrichment of categories corresponding to metabolic process and proteasomal protein catabolic process (Fig.3e). The GO analysis of CC (Cellular Component) found that the Ψ-containing transcripts were highly enriched (*p* < 2.0e-09) in the mitochondrial inner membrane. These results indicate that Ψ-containing transcripts are involved in cellular metabolism and could be important for mitochondria function (Supplementary Fig.3d).

Within mRNA, Ψs are distributed along the 5’ UTR, CDS, and 3’ UTR, with an enrichment in CDS and an underrepresentation in 5’ UTR (Fig.3f), consistent with our previous finding^6^. We then analyzed the codon preference of Ψ sites that are present in the single and double U contexts (accounting for approximately 39% and 40% of all Ψ sites) (Supplementary Fig.3e). We found that UUC and GUU codons, encoding phenylalanine and valine, are the most frequently modified (Fig.3g, Supplementary Fig.3f). Ψ is mainly present in the second and third position; interestingly, 2 Ψ sites were detected in the AUG start codons, while 2 Ψ sites were detected in UAG stop codons (Fig.3d, Supplementary Fig.3f).

Although transcriptome-wide Ψ distribution has been investigated previously^6,21–23^, the absolute level of Ψ modification in mRNA has not been reported. We first analyzed the modification level of the 2,174 Ψ sites, and found that their deletion rates are highly reproducible (R^2^ = 0.946, Fig.3h). Globally, ~80% of the Ψ sites were modified at low level (Ψ level < 20%) in HEK293T mRNA and ~10% of the sites were highly modified (Ψ level > 40%) (Fig. 3i). The median modification level of mRNA Ψ sites was about 10%, which is similar to the methylation level of m^5^C in human mRNA^45^.

We next analyzed the stoichiometry of Ψ sites in different sequence contexts. We found that 39% Ψ sites possess a modest “GUG” motif and that 40% “UU” sites prefer a “GUUCNA” motif (N=A/G/C/U) (Fig.3j). The top 10 Ψ motifs are shown in Fig 3k: “UGUAG” is the most enriched motif, resembling the preferred sequence context of PUS7, while the “GUUCNA” motif is reminiscent of the reported targets of TRUB1 (Fig.3k). Ψ sites conforming to the “GUUCNA” motif exhibit a much higher modification level (median Ψ level > 40%) than those within the “UGUAG” motif (5%~15% median level, Fig.3l). Thus, such distinct Ψ context and modification level suggests the differential activity of various sequence-specific modification machineries towards human mRNA.

### The quantitative, PUS-dependent Ψ maps

We next sought to define the molecular basis for targeting the mRNA and ncRNA transcripts for pseudouridylation. Previous efforts to map Ψ sites yielded sites that could be attributed to PUS1/PUS7/TRUB1^6,21,23,32^, yet the confidence and stoichiometry information is limited due to the inherent property of the CMC-based methods. To determine if PRAISE could precisely and quantitatively map PUS-dependent Ψ sites, we applied it to several HEK293T cell lines we generated previously^6,46^, with one candidate PUS enzyme been knocked out at a time. In addition, we generated a new TRUB1 KO HEK293T cell line; using quantitative MS, we found a notable decrease (~4%) of Ψ level for the small RNA (< 200 nt) population, consistent with a role of TRUB1 in modifying tRNA and thus the usefulness of the new TRUB1 KO cell line (Supplementary Fig.4a).

We applied PRAISE to the cellular polyA+ RNA of three KO cell lines. Dependent-Ψ sites were defined by comparing the Ψ profile between WT and KO. Overall, we identified the 37, 165 and 346 targets for PUS1, PUS7, and TRUB1, respectively (Fig.4a-c, Supplementary table 3). The median modification level of PUS1- and PUS7-dependent Ψs are ~10%, while TRUB1-dependent Ψ sites are highly-modified and their median level is ~35% (Fig. 4d-f), indicating a high enzymatic activity of TRUB1 *in vivo*. To understand the specificity of enzymes toward their targets, we examined the sequences and structural elements surrounding the PUS-dependent Ψ sites. TRUB1-dependent Ψ sites are highly enriched in a ‘‘GUUCNA” consensus sequence (Fig.4i), while The majority of PUS7-dependent Ψ sites are highly enriched in a ‘‘UGUAG” consensus sequence (Fig.4h) and PUS1 targets shared a weak “GUG” sequence motif (Fig.4g). The secondary structure of the 346 TRUB1 mRNA targets were predicted (see Methods), and they were predicted to give rise to a typical hairpin, consisting of a 5-bp stem and a 7-nt loop, with the Ψ site being the second base in the loop (Fig.4j). Thus, TRUB1 appears to recognize a conserved motif and hairpin structure in mRNA and tRNA. On the other hand, the structural constraint was modest for the PUS1 - and PUS7-dependent Ψs (Supplementary Fig.4e,f).

**Figure 4.**
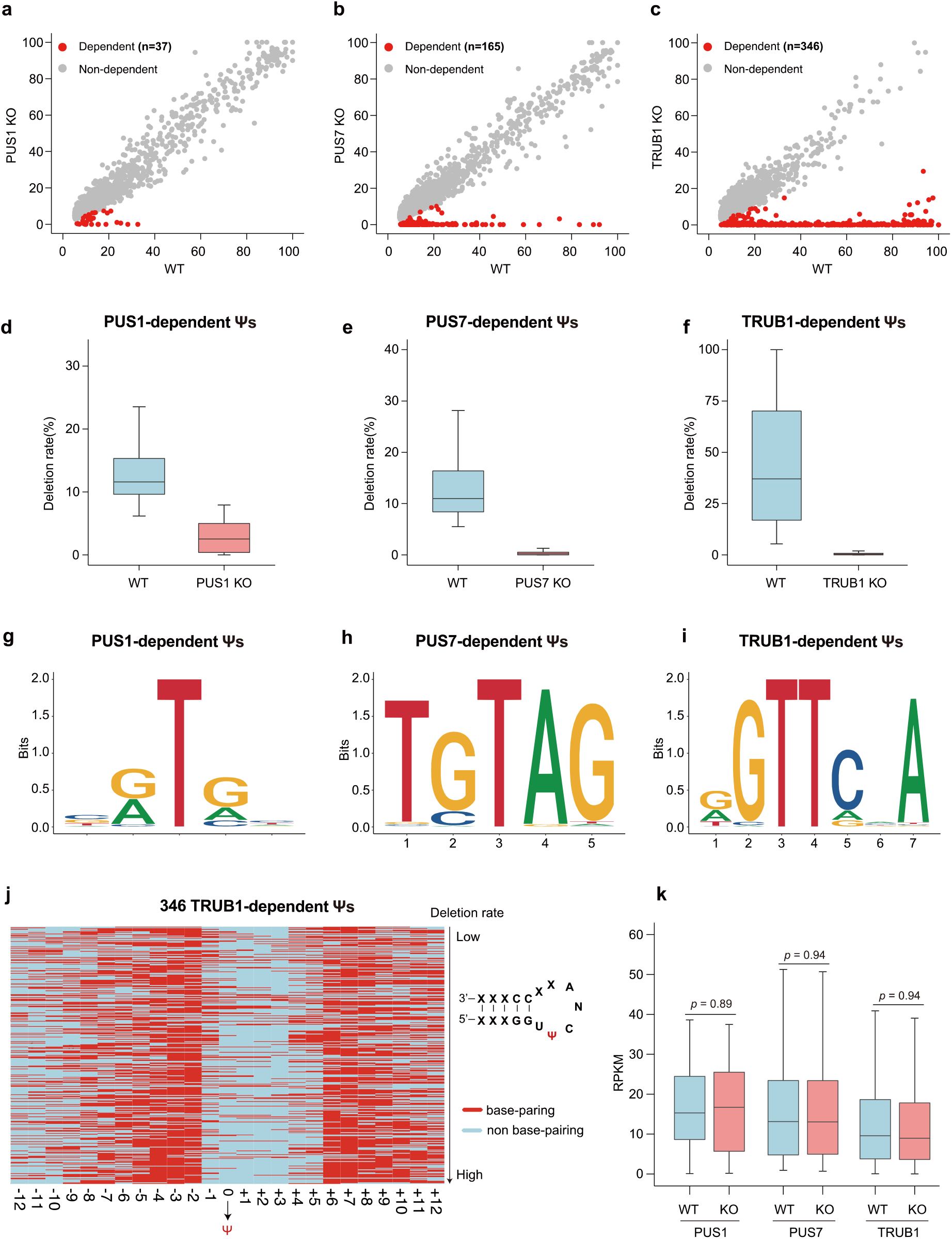
The quantitative, PUS-dependent Ψ maps. (a-c) Scatterplot of Ψ sites of WT versus PUS1/PUS7/TRUB1-KO cells. The red and gray dots denote the dependent Ψ sites and non-dependent Ψ sites, respectively. (d-f) Boxplot showing the deletion rate of the KO-dependent Ψ sites in WT versus PUS1/PUS7/TRUB1 KO treated samples. (g-i) Motif analysis of PUS1/PUS7/TRUB1-dependent Ψ sites identified by PRAISE. (j) Heat map depicting the secondary structure of 346 TRUB1-dependent Ψ sites. The red color denotes base pairing while blue color denotes non-base-paring. (k) Boxplot showing the RPKM level of dependent-Ψ transcripts in WT versus PUS1/PUS7/TRUB1 KO cells.

We also analyzed the distribution of the PUS-dependent Ψ sites. They generally resemble the overall distribution pattern of Ψ (Supplementary Fig.4b,c,d), with PUS7- and TRUB1-dependent sites more enriched in 3’UTR (Supplementary Fig.4c,d). We then examined potential role of Ψ in gene expression regulation. We compared the expression levels of transcripts containing PUS-dependent Ψ, and found no difference in expression level between the wild type and the corresponding KO cells (Fig.4k, Supplementary table 4). This observation suggested that Ψ does not affect RNA stability.

### Mitochondrial Ψ landscape

In addition to the transcriptome-wide Ψ modification in nuclear-coded transcripts, we also investigated Ψ sites in the mitochondrial transcripts. The mitochondrial genome encodes 2 mt-rRNAs, 22 mt-tRNAs, and 13 mt-mRNAs of the oxidative phosphorylation (OXPHOS) system. In total, we identified 2, 7 and 4 Ψ sites from rRNA, tRNA and mRNA in mitochondrial heavy strand, respectively (Figure 5a, Supplementary table 5). No Ψ site was found in mitochondrial light strand. The modification rate of the two mt-rRNA is 38.4% and 6.6% for Ψ3067 and Ψ1207-1208 respectively, which appears much lower than their rRNA counterpart. The highest modification site identified is mt-Leu tRNA(Ψ3286), showing a modification stoichiometry of 90.2%.

**Figure 5.**
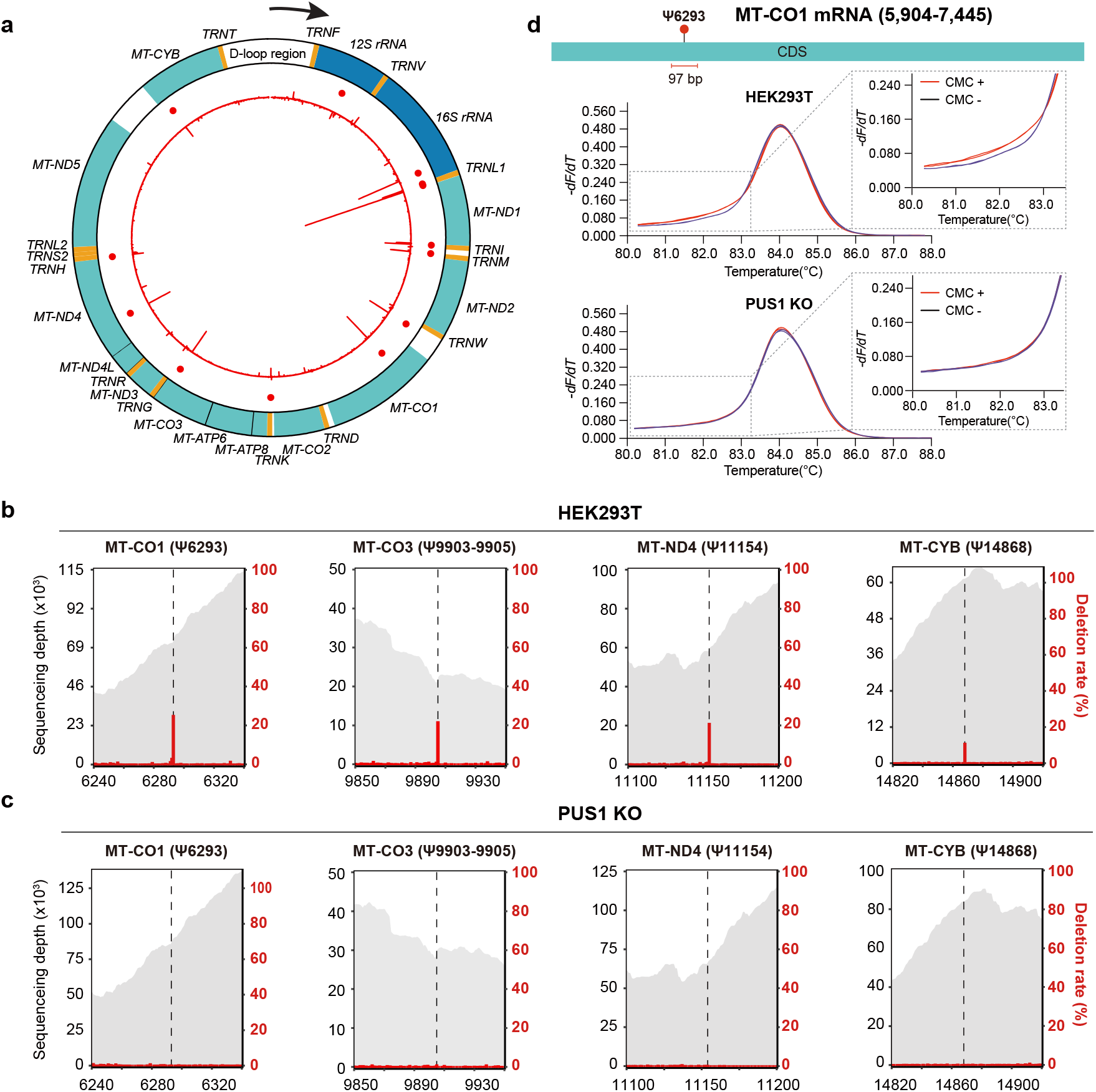
Mitochondrial Ψ landscape and Ψ enzymes. (a) Ψ sites identified in the heavy strand of mitochondria. Orange, blue and green colors represent tRNA, rRNA and mRNA, respectively. The red line in the inner circle represents the difference in deletion rate (treated-untreated) of individual nucleotides, while each red dot represents an identified Ψ sites. (b) The sequencing depth of regions surrounding four mt-mRNA Ψs and the corresponding deletion rate in HEK293T (WT) treated samples are plotted. (c) The sequencing depth of regions surrounding four mt-mRNA Ψs and the corresponding deletion rate in PUS1 KO treated samples are plotted. (d) High-resolution melting analysis (HRM) results for position Ψ6293 in MT-CO1 mRNA, from the HEK293T (WT) and PUS1 KO cell lines.

For mt-mRNA, the Ψ stoichiometry ranges from 10 to 26%. Ψ sites at position 6294 in MT-CO1 mRNA and position 9904–9906 in MT-CO3 mRNA are known^47^; expectedly, they demonstrated the highest modification level among mt-mRNAs, which are 26.0% and 22.1% respectively (Figure 5b). A novel Ψ sites were identified in MT-ND4 and MT-CYB mRNA respectively, which are Ψ11154 and Ψ14868. They bear a modification level of about 21.2% and 10.4%, respectively (Figure 5b).

### Mitochondrial Ψ enzymes

We next aimed to explore Ψ synthases responsible for pseudouridylation in mt-RNAs. It is known that one of the two PUS1 isoforms contains a mitochondrial targeting signal (MTS) in its N-terminal region (Supplementary Fig.5a) and has been shown to modify Ψ27 and Ψ28 in mt-tRNAs^48^; we thus examined whether PUS1 may have additional targets in the mitochondria. We first performed immunofluorescence staining of the two PUS1 isoforms in Hela cells, and expectedly we found that PUS1 isoform 1 was mainly located in the mitochondria while isoform 2 was abundant in the nucleus (Supplementary Fig.5b). We then compared the mitochondrial Ψ profile between the wild type and PUS1 KO cells. We found that modification signals for 1,5, and 4 Ψ sites in mt-rRNA, mt-tRNA and mt-mRNAs disappeared completely upon PUS1 knockout, suggesting that they are PUS1-dependent (Figure 5c, Supplementary Fig.5c). We also validated the presence and PUS1-dependency of Ψ6293 in MT-CO1 mRNA using an orthogonal, locus-specific detection method that we established previously^46^. The modification level, as estimated by melting curve alteration, was medium in the wild type 293T cells, consistent with its 25% Ψ stoichiometry by PRAISE (Figure 5d). Upon PUS1 knockout, the melting-curve alteration disappeared completely, which is in good agreement with our results via PRAISE (Figure 5d).

We next sought to identify the modification enzyme for the Ψ3286 site in mt-Leu tRNA. Since it shows a “GUUCAA” motif, we suspected that it could be a catalyzed by TRUB1. Expectedly, the deletion signal of Ψ3286 was almost absent in TRUB1 KO samples (Supplementary Fig.5d), suggesting that Ψ occurs at position 3286 and is TRUB1-dependent. As a comparison, after the nuclear-localized PUS7 was knocked out, no significantly changes of Ψ stoichiometry were observed in any mt-RNAs (Supplementary Fig.5e).

## Discussion

In this study, we developed a new base-resolution technology that enables quantitative Ψ detection. PRAISE utilizes Ψ-monobisulfite adducts induced deletion signal during cDNA synthesis, which allows for more accurate detection for Ψ compared to CMC-based methods. All reagents used by PRAISE are commercially available, circumventing the multiple-step chemical synthesis required for azide-derivatized CMC compound in our previous method^6^. We demonstrated good reproducibility between two biological replicates in PRAISE (~73%). PRAISE also shows excellent quantitative capabilities in rRNA and spike-in RNA. In addition, PRAISE can be readily adopted for locus-specific and quantitative interrogation of Ψ sites^46,49^. The detailed mechanism for the generation of deletion signal in RT remains unclear at the moment; but it is clearly influenced by the type of RTases and the condition of RT. Future elucidation of the mechanism may help further optimize the method for more biological contexts.

Epitranscriptomic technologies to detect multiple modifications at the single-molecule resolution is highly desired but remains to be established^50^. Since PRAISE is based on readthrough of RT and induces deletion signal of Ψ per cDNA strand, it could be combined with orthogonal chemistry to map additional RNA modification types simultaneously^45,51,52^. Potential crosstalk of diverse RNA modifications may be investigated via such approaches. For instance, m^6^A has been found to modulate A- to-I RNA editing^53^. In addition, third-generation sequencing technologies, such as Nanopore sequencing, have the potential to directly detect multiple modifications relying on various signatures including base-calling “errors”^29,31,54–56^; yet, they await further improvement in terms of accuracy and sensitivity. The various chemical/biochemical treatments developed for different modifications, including the PRAISE methodology reported herein, can be coupled to third-generation sequencing platforms to directly identify multiple modifications including Ψ at the single-molecule level.

Our results showed that Ψ is enriched in coding sequence of transcripts. Because it has a similar Watson-Crick interface with U, Ψ is not expected to directly alter the decoding of a sense codon. However, Ψ in stop codons has been found to suppress translation termination and promote protein readthrough^57^. We found rare but existing cases in which Ψ is present in the stop codon of endogenous mRNA transcripts (2 sites); it remains to be determined whether or not these Ψ sites mediate stop codon readthrough in vivo. Also, we found two examples in which the start codon is modified by Ψ; whether or not it may alter translation initiation is unclear. In addition, our analysis focused on Ψ distribution and level in mature mRNA; it is known that Ψ could be installed co-transcriptionally and function during mRNA processing, for instance, alternative pre-mRNA splicing^14^. Thus, PRAISE can be readily applied to reveal the modification level of Ψ in nascent transcripts. Moreover, an interesting question is whether Ψ is present in chromatin-associated regulatory RNA and might be involved in the regulation of chromatin state and gene expression, as recently demonstrated for m^6^A^58–61^.

PRAISE enabled us to not only unambiguously assign Ψ sites to certain PUSs, but also reveal their modification stoichiometry. Because human contains 13 PUS enzymes, it has been speculated that redundancy may exist in terms of mRNA pseudouridylation^21,62^. Yet, among PUS1, PUS7 and TRUB1, our data in 293T cells reveals very specific mRNA targets; the PUS-dependent Ψ sites do not overlap with each other. Future studies may examine whether or not PUSs belonging to the same family may have potential redundancy in modifying mRNA. Functionally, PUS7-mediated Ψ has been implicated previously to stabilize transcripts under heat shock conditions^23^; yet, our results comparing WT and PUS7 KO cells revealed no effect of Ψ on mRNA stability. This could be due to differential biological contexts of the two studies. Alternatively, it is tempting to speculate that Ψ could function in biological processes other than mRNA stability, for instance, in discriminating “self” from “non-self’ RNA in nucleic acid sensing^63^, which is an established role for inosine^64^. Indeed, incorporating Ψ and its methylation derivatives in foreign RNA allows for escape from innate immune sensing, which is believed to contribute to the major success of COVID-19 mRNA vaccines^9,10^. We hope that the transcriptome-wide, quantitative power of PRAISE could aid new discoveries of the ancient Ψ modification as well as future development of therapeutics involving RNA modifications.

## Methods

### Cell culture

HEK293T were cultured in DMEM medium (Corning) supplemented with 10% (v/v) FBS (Gibco), 1% GlutaMAX (Gibco) and 0.5% penicillin/streptomycin (Gibco) at 37 °C with 5% CO_2_. Cells within passage 3-6 were used for experiments.

### Generation of stable knockdown or CRISPR knockout cell lines

The shRNA targeting DKC1 (TRCN0000010325) was cloned into pLKO.1 vector. A scrambled shRNA was used as the mock control. Cell transfection was performed according to Broad Institute. The knockdown efficiency was verified by qPCR and western blot. Sequences of oligoes and qPCR primers of DKC1 were listed as follows: DKC1-qFWD: GATATGAGGACGGCATTGAG; DKC1-qRVS: GGTCGCAGGTAGAGATGA PUS1 KO HEK293T cells and PUS7 KO HEK293T cells were from the literature^6,46^. TRUB1 KO HEK293T cells were generated by CRISPR–Cas9 technology^65^. sgRNA sequences of TRUB1 was listed as follows: TRUB1-sgRNA: TTCGGATCCGGTCCTGGCCG

### mRNA purification

Total RNA was extracted with TRIzol (Life technologies) followed by isopropanol precipitation, according to the manufacturer’s instructions (Invitrogen). The resulting total RNA was treated with DNase I (NEB) to avoid DNA contamination. For PolyA+ RNA isolation, size selection was first performed using MEGAclear Transcription Clean-Up Kit (Ambion) to deplete small RNA, and RNA was subsequently purified with two sequential rounds of polyA tail purification using oligo(dT)_25_ Dynabeads (Invitrogen).

### Quantification of Ψ level by LC-MS/MS

200 ng isolated mRNA was digested into single nucleosides by 0.5 U nuclease P1 (Sigma, N8630) in 20 μl buffer containing 10 mM ammonium acetate, pH 5.3 at 42 °C for 6 h, then mixed with 2.5 μl 0.5 M MES buffer, pH 6.5 and 0.5 U Shrimp Alkaline Phosphatase (NEB, M0371S), in a final reaction volume of 25 μl adjusted with water, and incubated at 37 °C overnight. The nucleosides were separated by ultraperformance liquid chromatography with a ZORBAX SB-Aq column (Agilent), and then detected by triple-quadrupole mass spectrometer (AB SCIEX QTRAP 6500). A multiple reaction monitoring (MRM) mode was adopted: *m/z* 245.0 to 179.1 for Ψ, m/z 245.0 to 113.1 for U. 5 μl of the solution was injected into LC-MS/MS. Standard curves were generated by running a concentration series of pure commercial nucleosides (Berry associated). Concentrations of nucleosides in RNA samples were calibrated by standard curves.

### Synthesis of spike-in RNA oligoes

Five pairs of 150 nt synthetic RNA oligoes containing either Ψ or U were used to examine the deletion signal of Ψ in a quantitative manner. Short Ψ/U-oligoes (70 nt) and doner RNA oligoes (80 nt) for subsequent ligation were synthesized by Shanghai Primerna Biotechnology (Shanghai, China). The 5’-end of the doner RNA were phosphorylated and 3’-end of the doner RNA were blocked by a ddC (2’,3’-deoxyCytosine) to avoid self-ligation. Splint ligation was performed according to the literature^66^. Briefly, a mixture of short Ψ/U-oligoes, doner RNA and the complementary splint DNA strand were annealed at a molar ratio of 1: 2: 1.5 in T4 DNA ligase buffer (NEB) by incubation for 3 min at 65°C, followed by 5 min at 25°C. Next, T4 RNA ligase 2 (NEB) and RiboLock RNase inhibitor (ThermoFisher Scientific) were added to the annealed mixtures and incubate at 37 °C for 1 h. Then the splint Ψ/U-oligoes were gel purified, selected regions of the gel corresponding to 150 nt were excised and recovered. Mixed Ψ-oligoes with paired U-oligoes at indicated ratios: 100%, 70%, 50%, 40%, 30%, 20%, 10%, 5%, 0%. 400 pg spike-in oligoes was mixed with 500 ng total RNA, and the Ψ stoichiometries were detected by PRAISE.

Splinted RNA oligoes sequences were listed as follows:

Oligo-0:

GUUGAUCUUGGCACGUCUACUCGACCGUCUGGAACUUGAUAUUAUCGGAUCAGUGAAUCACUCCAAGCCAAAUGGGCGGUAGGCGUGUACGGUGGGAGGUCUAUAUAAGCAGAGCUCUCUGGCUAACUAGAGAACCCACUGCUUACUGGC

Oligo-1:

AUCUACCUGUCCAGUAGCCUUCAGGAUCAUGCUGUCUGACUUGCUG**<u>U</u>**AGAU CAUCUAGUGCCAUAAGCCAAAUGGGCGGUAGGCGUGUACGGUGGGAGGUCUAUAUAAGCAGAGCUCUCUGGCUAACUAGAGAACCCACUGCUUACUGGC

Oligo-2:

ACGGUAAACUGCCCACUUGGCAGUACAUCAAGUGUAUCAUAUGCCAUG**<u>U</u>**AGG CCCCCUAUUGACGUCAAUCUUGAGUGGAGAGGCUAUUCGGCUAUGACUGAGCACAACAGACAAUCGGCUGCUCUGAUGUCGCUGUGUUACGGCUGUCA

Oligo-3:

GGCAUUAUGCCCAGUACAUGACCUUAUGGGACUUUCCUACUUGGCUG**<u>U</u>**AGA UCUACGUAUUAGUCAUCGCCUUGAGUGGAGAGGCUAUUCGGCUAUGACUGAGCACAACAGACAAUCGGCUGCUCUGAUGUCGCUGUGUUACGGCUGUCA

Oligo-4:

AGCUGGCUAGCGUUUAAACUUAAGCUUGGUACCGAGCUCGGAUCCUG**U**AGU CCAGUGUGGUGGAAUUCUGCUUGAGUGGAGAGGCUAUUCGGCUAUGACUGAGCACAACAGACAAUCGGCUGCUCUGAUGUCGCUGUGUUACGGCUGUCA

### Locus detection of Ψ1367 in 18S rRNA for bisulfite/sulfite conditions screening

#### RNA preparation

500 ng total RNA for one sample was treated with DNase I at 37°C for 30 min, then RNA was fragmentated to ~150 nt by magnesium RNA fragmentation buffer (New England Biolabs) for 4 min at 94°C, followed by chilling on ice. Reaction was stopped by RNA fragmentation stop solution and purified by ethanol precipitation. After ethanol precipitation, RNA was dissolved in 5 μl nuclease-free water.

#### Bisulfite/sulfite treatment

Standard bisulfite treatment referred to the conditions of literatures^36,37^, the bisulfite solution was freshly prepared by dissolving 4.05 g of sodium bisulfite (Sigma Aldrich) in 5.5 ml of RNase-free water, adjusting the pH to 5.1 with 10 M sodium hydroxide, and adjusting the volume to 10 ml with water. The 100 mM hydroquinone was prepared freshly by adding 11.01 mg of hydroquinone (Sigma Aldrich) to 1 ml of RNase-free water. For standard bisulfite treatment, 5 μl of RNA fragments was dissolved in 50 μl bisulfite solution, which is a 100:1 mixture of bisulfite solution and 100 mM hydroquinone, and subjected to heat incubation at 50°C for 16 h.

For our improved bisulfite/sulfite treatment conditions, weighed and mixed potassium sulfite and sodium bisulfite in the proportions. The 100 mM hydroquinone was prepared freshly by adding 11.01 mg of hydroquinone (Sigma Aldrich) to 1 ml of RNase-free water. For bisulfite/sulfite treatment, 5 μl of RNA fragments was dissolved in 50 μl bisulfite/sulfite solution, which is a 100:1 mixture of bisulfite/sulfite solution and 100 mM hydroquinone, and subjected to heat incubation at 70°C for 5 h. The reaction mixture was desalted by twice passing through Micro Bio-spin 6 chromatography columns (Bio-Rad).

#### Desulfonation

Desalted RNA was transferred to a new 1.5 ml Nuclease-free tube and adjusted to 100 μl with RNase-free water, then incubated with an equal volume of 1 M Tris-HCl (pH 9.0) at 75°C for 30 min. The reaction was then immediately stopped by chilling on ice and ethanol precipitation.

#### RT-PCR

bisulfite/sulfite treated RNA was reverse transcribed into cDNA using random hexamers (Thermo Fisher) with Maxima H minus Reverse Transcriptase (Thermo Fisher) according to the manufacturer’s instructions. Briefly, after ethanol precipitation, the RNA was resuspended in 10 μl of RNase-free water, with the addition of 1 μl 100 μM RT primer. RNA-primer mix was denatured at 80°C for 2 min followed by chilling on ice immediately. 4 μl 5X First-strand buffer, 1 μl 10 mM dNTPs, 2 μl 0.1 M DTT, 1 μl 40 U/μl RNase Inhibitor and 1 μl Maxima H minus were added into the denatured RNA-primer mix. Reverse transcription was performed by incubating at 25°C for 5 min, 42°C for 3 h and heat-inactivated at 70°C for 15 min. Next, 1 μl of the cDNA was used for PCR reaction (Total volume of 50 μl) using specific forward and reverse primer of Ψ1367_18S and Zymotaq DNA polymerase for 34 cycles. 5 μl PCR products for Ψ1367_18S were then assessed on 3% agarose gel. The remaining PCR products were recovered, cloned into TOPO-TA cloning vectors and transformed to E. coli competent cells. The deletion rate and C-to-T conversion rate were assessed by Sanger sequencing at least 25 individual clones. PCR primer of Ψ1367_18S were listed as follows:

Ψ1367_18S-FWD: AGGGCACCACCAGGAGTGGAGCC
Ψ1367_18S-RVS: TAACTAGTTAGCATGCCAGAGTCT

### Comparison of different reverse transcriptases

Commercially available reverse transcriptases were compared to determine which enzyme caused highest deletion rate and yielded the most cDNA. Ten reverse transcriptases including Maxima H minus (Thermo Fisher), SuperScript II (Thermo Fisher), SuperScript III (Thermo Fisher), SuperScript IV (Thermo Fisher), Revert Aid (Thermo Fisher), M-MLV (Thermo Fisher), AMV (New England Biolabs), ProtoScript II (New England Biolabs), Recombinant HIV (Worthington) and HiScript III (Vazyme) were tested. All RT enzymes were used according to the manufacturer’s protocol. The yield in treated samples of AMV and Revert Aid were too low to be used for sequencing and analysis. For another eight reverse transcriptases, the deletion rate of all rRNA Ψ sites was compared.

### PRAISE

#### Fragmentation

First, 500 ng DNase I treated total RNA or mRNA of one technical replicate was fragmentated to ~150 nt by magnesium RNA fragmentation buffer (New England Biolabs) for 4 min at 94°C, followed by chilling on ice. Reaction was stopped by RNA fragmentation stop solution and purified by ethanol precipitation. After ethanol precipitation, RNA was resuspended in 6 μl nuclease-free water. 1 μl of fragments (~75 ng) are transferred into a new nuclease-free tube and used as “untreated” and stored at −80°C. Remained RNA was subjected to bisulfite/sulfite treatment.

#### Sulfite/bisulfite treatment

The 85% sulfite/15% bisulfite solution was prepared freshly by adding 2.58 g of potassium sulfite (Sigma Aldrich) and 0.30 g sodium bisulfite (Sigma Aldrich) to 8 ml of RNase-free water and the solution was vortexed to clear (pH value is ~7.5). The 100 mM hydroquinone was prepared freshly by adding 11.01 mg of hydroquinone (Sigma Aldrich) to 1 ml of RNase-free water. For sulfite/bisulfite treatment, 5 μl of RNA fragments was dissolved in 50 μl sulfite/bisulfite solution, which is a 100:1 mixture of 85% sulfite/15% bisulfite solution and 100 mM hydroquinone, and subjected to heat incubation at 70°C for 5 h. The reaction mixture was desalted by twice passing through Micro Bio-spin 6 chromatography columns (Bio-Rad).

#### Desulfonation

Desalted RNA was transferred to a new 1.5 ml Nuclease-free tube and adjusted to 100 μl with RNase-free water, then incubated with an equal volume of 1 M Tris-HCl (pH 9.0) at 75°C for 30 min. The reaction was then immediately stopped by chilling on ice and ethanol precipitation. After ethanol precipitation, the RNA was resuspended in 10 μl of RNase-free water, and the concentration of RNA was quantified by Qubit RNA HS Assay Kit (Thermo Fisher Scientific, Q32855). Take 50 ng RNA (~1 μl) as “treated” sample for subsequent library construction.

#### Library construction

50 ng “treated” RNA sample and 10 ng “untreated” RNA sample were subjected to library construction using SMARTer^®^ Stranded Total RNA-Seq Kit v3 - Pico Input Mammalian (Takara Bio, 634485) according to the manufacturer’s protocol with several modifications. Briefly, for first-strand cDNA synthesis, Maxima H minus Reverse Transcriptase (Thermo Fisher) was used instead of SMARTScribe II Reverse Transcriptase. Second, pre-PCR library was purified twice by 0.8X AMPure^®^ XP reagent and finally eluted by nuclease-free water. Since Ψ-containing rRNA fragments in sulfite/bisulfite treated sample can induce deletion signal, it cannot be digested by R-Probes v3, thereby potentially interfering with quantification of Ψ sites in rRNA. Therefore, we note that the step of ribosomal cDNA depletion was skipped. 11 cycles were used for untreated and treated RNA in the second round of PCR. Remained procedures were performed with the SMARTer Stranded Total RNA-Seq Kit v3 (Pico Input Mammalian) according to the manufacturer’s instructions. Library sequencing was performed on llumina Hiseq X10 with paired-end 2X150 bp read length.

### Pre-processing of raw sequencing data of PRAISE

Strand orientation of the original RNA as preserved on the process of library construction and reads R2 yields sequences sense to the original RNA. Thus, only reads R2 was used in our study. Illumina sequencing reads were first treated with cutadapt (version 3.5) for adaptor removal and quality trimming. Work command are as follows: cutadapt -j 28 --times 1 -e 0.1 -O 3 --quality-cutoff 25 -m 55. We then used Seqkit (version 0.13.2) to deduplicate PCR of Takara library based on 8 bp UMI at the 5’ end of reads R2, key process parameters are as follows: seqkit rmdup -s. Finally, umi_tools (version) was used to remove the 8bp UMI in the deduplication read. Six bases after the UMI added during library construction at the 5’ end of inserted sequences and the six bases in at the 3’ end of inserted sequences was also removed, using the key parameters: umi_tools extract --extract-method=string --bc-pattern=NNNNNNNNNNNNNN and umi_tools extract --extract-method=string -- 3prime --bc-pattern=NNNNNN.

### PRAISE-tools and code availability

After the reads cleaning steps, analysis of PRAISE data mainly includes two major parts, which is alignment of reads and identification of pseudouridine sites with quantified deletion signal level. To make the analysis pipeline easy to implement, we developed PRAISE-tools, a computational pipeline that quantifies the deletion signals with high confidence. PRAISE-tools takes cleaned reads as input and finally reports the deletion rate of T sites as modification level.

### Alignment of reads

PRAISE-tools firstly got all possible mapped position to reference transcriptome (GRCh38) using HISAT2 (version 2.2.1) very sensitive mode. The key parameters are as follows: hisat2 -q --repeat --no-spliced-alignment --very-sensitive. After removing unmapped reads, using our own tailored realignment that based on the customized biopython pairwise alignment to remove low confidence alignments and get accurate deletion sites. The key parameter is python realignment.py –fast -ms 4.8. Next, the multiple-mapping reads with lower mapping score will be removed and the deletion signal just one bp before reads’ terminal will be removed using remove_multi-mapping.py and remove_end_signal.py. After getting the final mapping results in BAM, convert it into mpileup format with the key parameter of samtools mpileup -d 1500000 -BQ0 --ff UNMAP, QCFAIL -aa. Finally, the mpileup file will be counted and transferred into a specific BMAT format file.

### Call Ψ signal

To call Ψ signal, we next applied statistical test to each of T sites (contingency table test between treated sample and untreated sample). The calculated significance of p value is required to be <0.0001, the calculated deletion rate difference between treated and untreated sample is required to be > 5%. Since we mapped read to the transcriptome, the same sites will be counted repeatedly in the different transcripts of one gene. Therefore, we finally remove those repeated signals in one gene by converted sites in the transcript into sites in the genome, and kept the longest transcript using site_merge.py to give the Ψ lists of one sample. To give our final Ψ signal list, we overlapped two lists from two samples, only counted overlapped signals into our final Ψ list, and take the average deletion rate of two samples as its Ψ level.

### Call Ψ signal in KO samples

To call Ψ signal in KO samples, we took our final Ψ list as reference to find the deletion rate in KO samples. Then, we calculate the deletion rate difference between KO samples and WT samples as well as the deletion rate decreasing rate. For Enzyme dependent sites, the deletion rate difference is required to > 5%, and the decreasing rate of deletion is required to > 5%. To exclude the situations that the sites are not covered in the KO samples, the total counts in KO sample is required to 10. To give our final KO dependent sites list, we overlapped two lists from two KO repeats, only counted overlapped signals into our final KO dependent list.

### RNA secondary structure prediction

To analysis RNA secondary structure, we first extract the sequences near the Ψ sites (±12 nt), sort them by the deletion rate of Ψ to create a fasta file. Then, we used RNAfold to predict RNA secondary structure with following key parameters, RNAfold--temp=37 -p.

### Quantification and statistical analysis

The p-values were calculated using unpaired two-sided Mann-Whitney U-test. *\*P* < 0.05; \*\**P* < 0.01; \*\*\**P* < 0.001; ns, not significant. Error bars represent mean ± SD.

## Data and software availability

Sequencing data have been deposited into the Gene Expression Omnibus (GEO) under the accession number: GSE212210.

## Acknowledgements

The authors would like to thank Prof. Guanzheng Luo for providing IVT RNA; Drs. Haowei Meng and Bo He for advice on tailored alignment, Ms. Ruilin Xu for help with TA cloning experiments; Drs. Xushen Xiong and Mr. Hao Wu for advice on mitochondrial analysis; Dr. Jinghui Song and Ms. Hanxiao Sun for discussions. We thank National Center for Protein Sciences at Peking University for assistance with fragments analysis and NGS experiments; High Performance Computing Platform of the Center for Life Science for assistance with the bioinformatic analysis. This work was supported by the National Key Research and Development Program of China (nos. 2019YFA0110900 to C.Y., 2019YFA0802200 to C.Y., and 2021YFC2302400 to M.Z.), National Natural Science Foundation of China (nos. 21825701 to C.Y. and 92153303 to C.Y.), and China Postdoctoral Science Foundation (2020M680219 to M.Z.).

## Author contributions

M.Z., Z.J. and C.Y. conceived the project and wrote the manuscript. M.Z. and Z.J. developed the chemical assay under the guidance of C.Y., M.Z. designed and performed the experiments with the help of Y.M., Z.J. designed the tailored alignment method and performed the bioinformatic analysis of NGS data with the help of W.L. and K.L., Y.Z. performed orthogonal locus-specific experiments and immunofluorescences. B.L. constructed TRUB1 knockout cells and helped screen RT conditions, C.Y. supervised the project.

