## Supplementary information for "Quantitative profiling of pseudouridylation landscape in the human transcriptome"

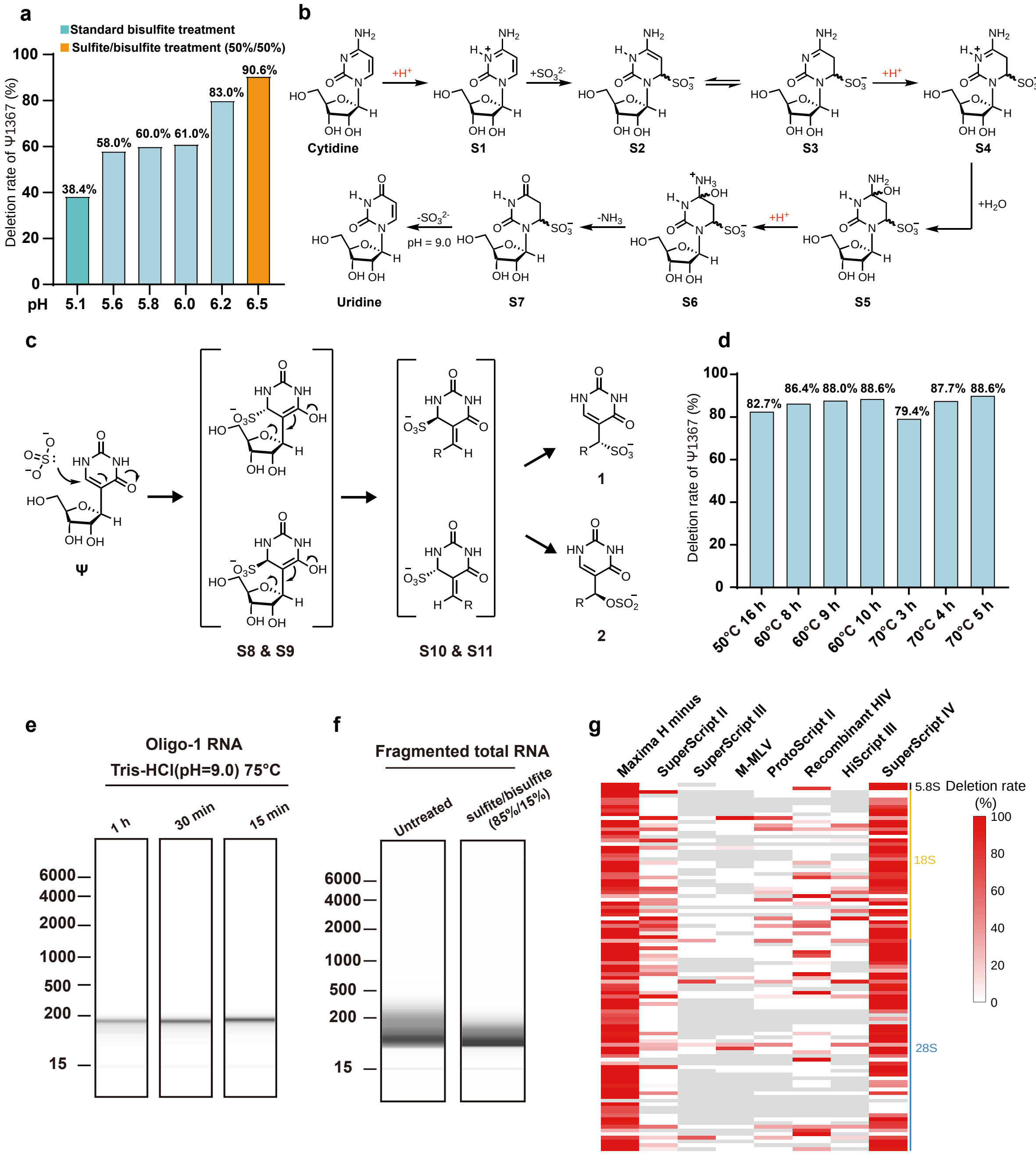


**Figure S1 | Performance of different chemical reaction and RTase on capturing Ψ-induced deletion signature under different conditions.**

(a) The deletion rate of Ψ1367 site in 18S rRNA under different sulfite/bisulfite conditions. The proportion of sulfite and bisulfite was increased from 95% sulfite/5% bisulfite (named "Standard bisulfite treatment") to 50% sulfite/50% bisulfite ("50%

sulfite/50% bisulfite treatment"), and the pH values were showed in the horizontal axis. 50% sulfite/50% bisulfite reagent corresponding to pH 6.5.

(b) Chemical mechanism of conversion from cytidine to uridine. The process is both acid-dependent and sulfite-dependent.

(c) Chemical mechanism of reaction between pseudouridine and sulfite reagent. After Michael addition, Ψ-SO3 generates two isoforms. Then, after two steps of reverse Michael addition and Michael addition, the final product is two ring-open isoforms.

(d) The deletion rate of Ψ1367 site in 18S rRNA under different time and temperature conditions. The horizontal axis represents time and temperature, and the vertical axis represents the deletion rate.

(e) Fragments analysis of oligo RNA after Tris-HCl treatment for different time. "1h", "30min" and "15min" represent different treatment time.

(f) Fragment analysis of fragmented total RNA. "untreated" represents fragmented total RNA with no treatment and "sulfite/bisulfite (85%/15%)" represents fragmented total RNA with sulfite/bisulfite treated.

(g) Heatmap of deletion rate of Ψs in rRNA under different reverse transcriptase conditions.

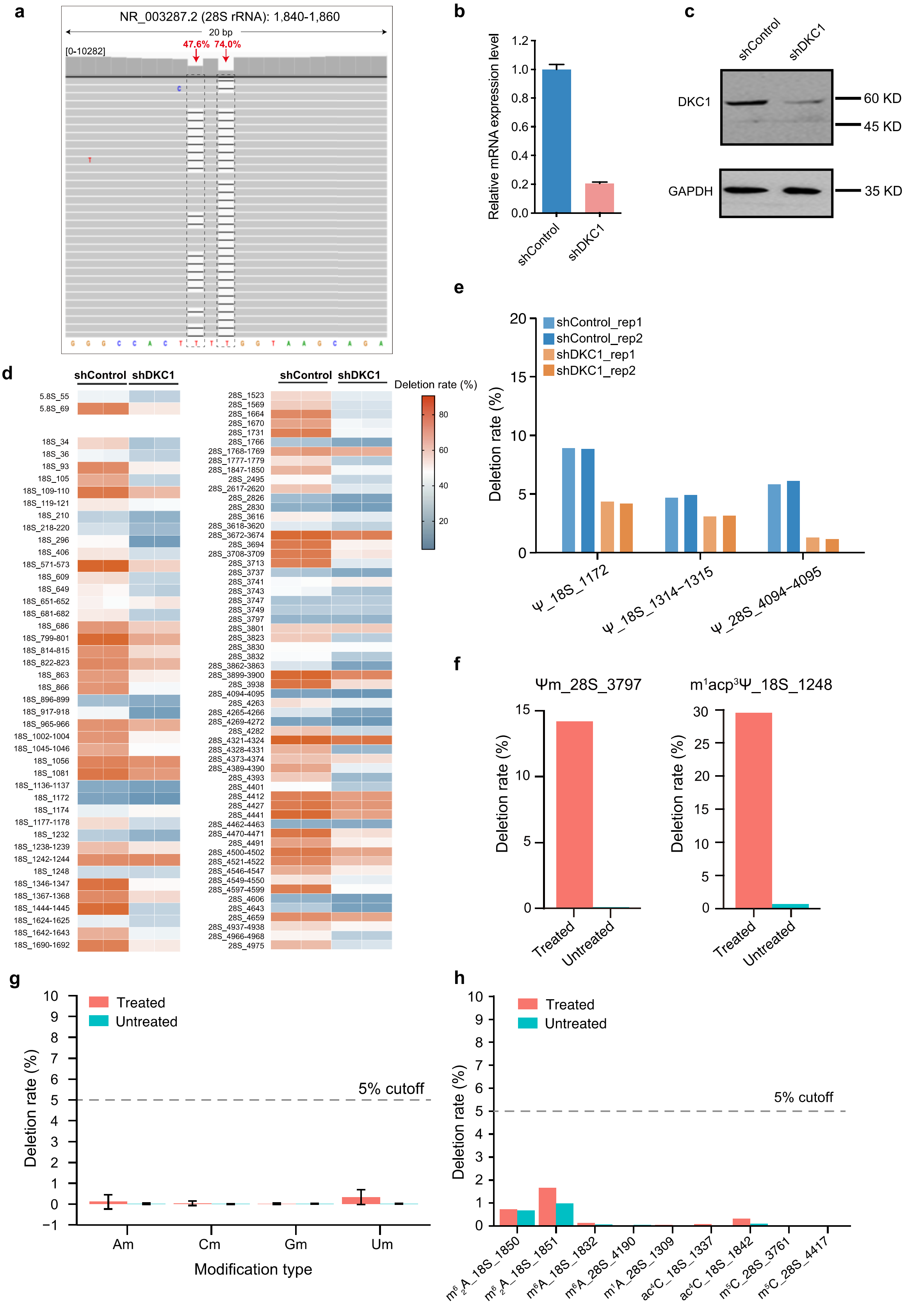


**Figure S2 | Identification of known and novel rRNA sites under DKC1 knockdown and evaluation of the deletion rate for other RNA modifications.**

(a) An IGV view of read mapping showing deletion signals in position 1840-1860 of 28S rRNA. Two deletion sites were marked between four consecutive Ts in 1847-1850.

(b) Boxplot representing the relative mRNA expression level of shControl and shDKC1 sample detected by qPCR. Data are mean ± SD; n = 2.

(c) The protein expression level of DKC1 knock-down cells was determined by Western blot. GAPDH was used as a loading control.

(d) Heatmap of deletion rate of all Ψ sites in 5.8S rRNA, 18S rRNA and 28 rRNA between shControl and shDKC1 sample.

(e) Deletion rate of three additional Ψ sites in rRNA in shControl and shDKC1 sample. Each sites have two replicates of shControl and shDKC1 sample.

(f) Deletion rate of known Ψm and $m^1acp^3\Psi$ sites in rRNA between treated and untreated samples.

(g) Deletion rate of known Am, Cm, Gm, and Um sites in rRNA between treated and untreated samples.

(h) Deletion rate of known $m^6_2A$, $m^6A$, $m^1A$, $ac^4C$, and $m^5C$ sites in rRNA between treated and untreated samples.

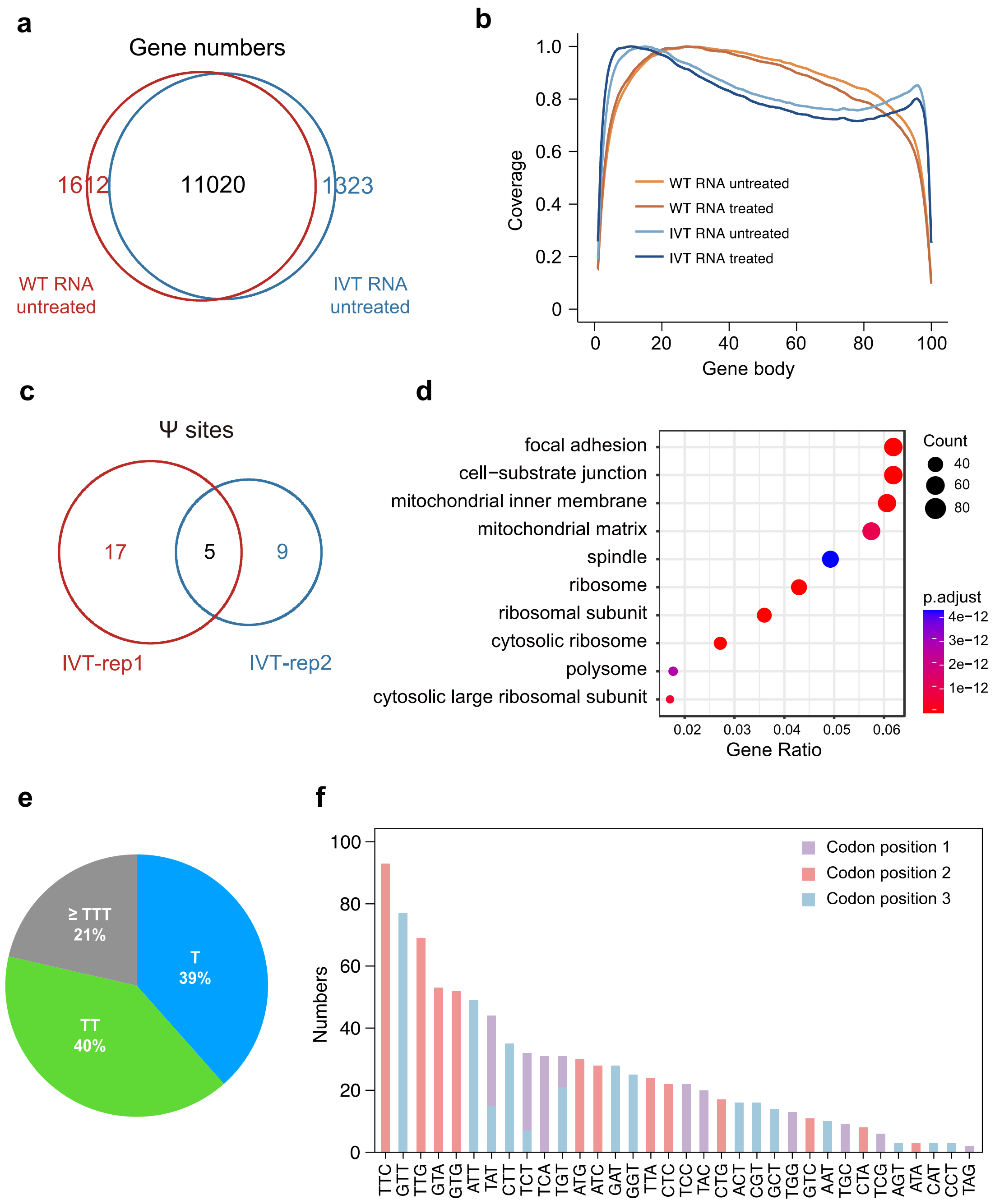


**Figure S3 | Quality control of IVT samples and codon usage of Ψ sites.**

(a) Venn diagram depicting gene overlap between in wild type cellular RNA sample (red) and IVT sample (blue). IVT, in vitro transcription.

(b) Read coverage from the 5’ end to the 3’ end of transcripts in wild type cellular RNA (orange) and IVT RNA (blue) libraries. Each library has two replicates and are combined and scaled to a normalized gene model.

(c) Venn diagram depicting overlap of Ψ sites between in IVT replicate 1 (red) and IVT replicate 2 (blue).

(d) Genes containing Ψ sites are analyzed using gene ontology of cellular components.

(e) Pie charts representing proportions of Ψ sites within “T”, “TT” and “⩾TTT” contexts in human mRNA.

(f) The counts of Ψ sites in codon types as well as positions are shown. Ψ sites observed at the first (purple), second (red) and third (blue) codon position are indicated.

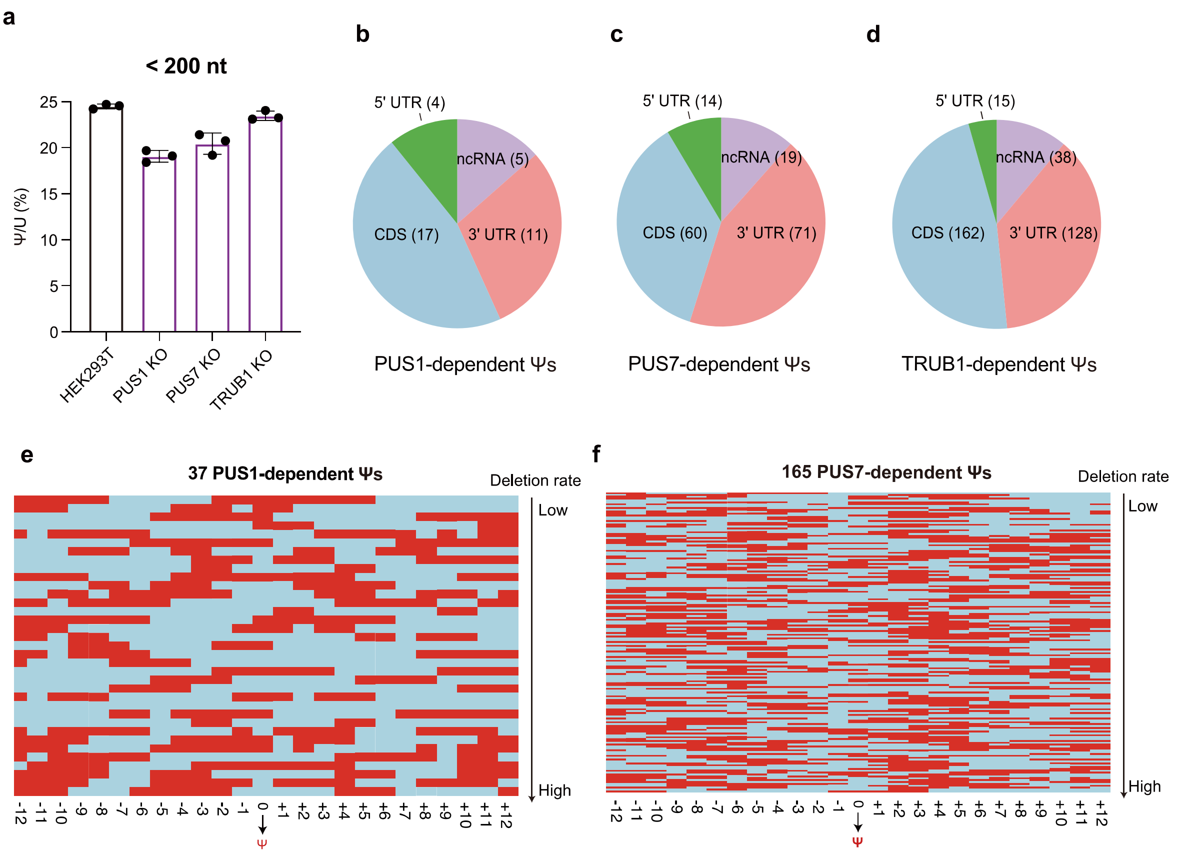


**Figure S4 | Distribution and secondary structure of KO-dependent Ψ sites.**

(a) Quantification of Ψ modification of small RNA (< 200nt) in wild type and PUS knocked out cell lines.

(b) Distribution of PUS1 dependent Ψ sites in human mRNA (32) and ncRNA (5). CDS, coding sequence.

(c) Distribution of PUS7 dependent Ψ sites in human mRNA (145) and ncRNA (19).

(d) Distribution of TRUB1 dependent Ψ sites in human mRNA (305) and ncRNA (38).

(e) Heatmap depicting secondary structure of 37 PUS1 dependent Ψ sites. A 24 nucleotides interval around Ψ site (at position 0) is selected. Paired bases are marked in red and unpaired bases are marked in blue.

(f) Heatmap depicting secondary structure of 165 RNA strands containing PUS7 dependent Ψ sites. A 24 nucleotides interval around Ψ site (at position 0) is selected. Paired bases are marked in red and unpaired bases are marked in blue.

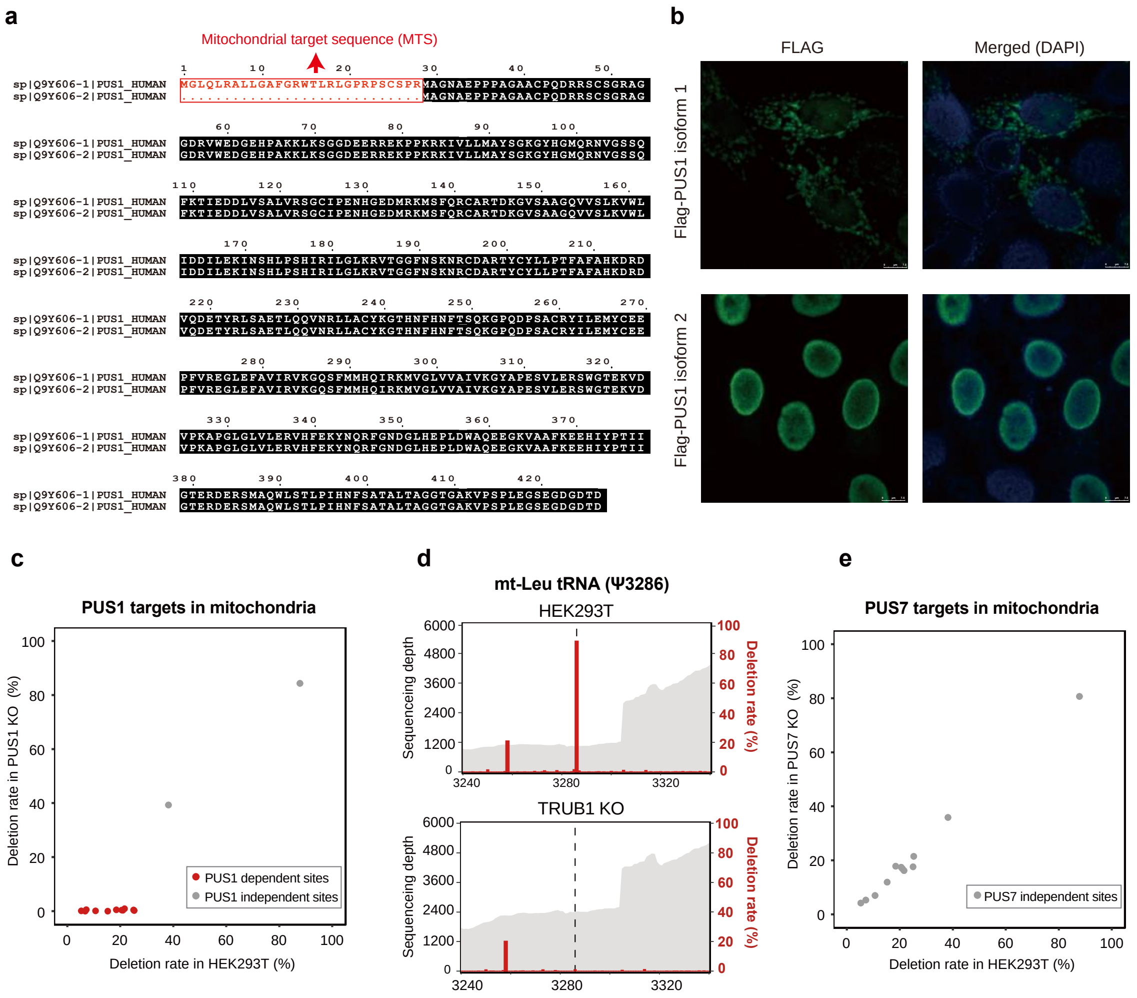


**Figure S5 | Subcellular localization of two isoforms of PUS1 and identification of mitochondrial targets assigned to PUS1 and PUS7.**

(a) Sequence alignment and comparation of PUS1 isoform 1 and isoform 2 in human cell line. Mitochondrial target sequence of PUS1 isoform 1 is marked in red color.

(b) Double immunofluorescence staining of HeLa cells transfected with expression vectors encoding Flag-tagged PUS1 isoform 1 and 2 (green) and DAPI (blue). The left panels show the expression vectors, and the right panels show the green and blue images merged in Adobe Photoshop.

(c) The deletion rate of Ψ sites in mt-RNA in PUS1 KO and HEK293T cells. The horizontal axis represents deletion rate of mt-RNA Ψ sites in HEK293T cells, and the vertical axis represents deletion rate of mt-RNA Ψ sites in PUS1 KO cells.

(d) The sequencing depth of regions surrounding Ψ3286 in mt-leu tRNA and the corresponding deletion rate in HEK293T (WT) and TRUB1 KO treated samples are plotted.

(e) The deletion rate of Ψ sites in mt-RNA in PUS7 KO and HEK293T cells.
